# Synthetic lethality between toxic amino acids, retrograde target genes and chaperones in *Saccharomyces cerevisiae*

**DOI:** 10.1101/2022.02.18.481112

**Authors:** Marina E. Druseikis, Shay Covo

## Abstract

The toxicity of non-proteinogenic amino acids has been known for decades. Numerous reports describe their antimicrobial/anticancer potential. However, these molecules are often toxic to the host as well, therefore a synthetic lethality approach can be beneficial. Here we investigated the potential synthetic lethality between toxic amino acids, the retrograde pathway, and molecular chaperones. In *Saccharomyces cerevisiae*, mitochondrial retrograde (RTG) pathway activation induces transcription of RTG-target genes, which replenishes alpha- ketoglutarate and glutamate; both metabolites are required for arginine and lysine biosynthesis. We previously reported that tolerance of canavanine, a toxic arginine derivative, requires an intact RTG pathway, and low-dose canavanine exposure reduces the expression of RTG-target gene*s*. At higher concentrations, canavanine causes protein misfolding. Here we show that in WT, low-dose thialysine exposure, a toxic lysine analog, has a similar effect as canavanine on RTG-target gene expression. To study if single amino acid deficiency and mild protein misfolding stress elicit a similar effect on RTG-target genes, we compared expression of heat shock protein (HSP) mutants grown without arginine or lysine to WT. Arginine deprivation induces RTG-target gene expression in *sse2Δ, hsp78Δ,* and *mdj1Δ*, but *CIT2* and *DLD3* are reduced in *cpr7Δ*. Lysine deprivation has the opposite effect on RTG-target gene expression in HSP mutants. Interestingly, exposure of HSP mutants to canavanine and thialysine reversed the trend of RTG-target gene expression in several cases. The RTG-target expression pattern in HSP mutants had a predictive value in canavanine sensitivity - the mutant with the lowest expression was the most sensitive - but this was not the case for thialysine. Some, but not all, mutants in RTG-target genes are sensitive to canavanine and thialysine; additional mutation in a certain HSP can exacerbate this sensitivity. Overall, we show that inhibiting molecular chaperones, RTG-target genes, or both can sensitize cells to low doses of toxic amino acids.

## Introduction

Canavanine, an amino acid analog of arginine, occurs naturally in leguminous plants. At high concentrations (20 - 200 µg/ml in *Saccharomyces cerevisiae*) [1–3] it causes protein misfolding and aggregation when incorporated into nascent proteins during translation. In this way, canavanine has antimicrobial properties and offers protection for the seeds containing it [4]. For years canavanine has been considered as a possible anticancer agent [5, 6], and recently has been shown to have potential as an anti-glioblastoma metabolic therapy when combined with arginine deprivation [7]. In fact, arginine deprivation alone has been shown to induce endoplasmic reticulum stress in solid human cancer cells [8] and arginine deprivation therapy (ADT) is currently being explored as a new metabolic targeting approach with high therapeutic potential for various solid cancers [9, 10].

We previously found that in yeast, certain genetic backgrounds and physiological conditions elicit major sensitivity to sublethal doses of canavanine (<1 µg/ml) [11]. Measuring proteotoxic stress at such low concentrations has proved challenging, since sublethal doses may not generate a strong enough response to meet the minimum threshold of detection [12] although the incorporation of canavanine into proteins can be detected at a media concentration of about 10 µg/ml [7]. We have shown that respiration-deficient yeasts (petites) produce less arginine than the respiratory WT and that canavanine induces mistargeting of arginine biosynthesis enzymes. Both factors contribute to canavanine sensitivity in petites [11].

Alongside petites, we found that strains defective in RTG genes (*rtgΔ* mutants) are highly sensitive to canavanine when grown on glucose lacking arginine (-ARG). RTG genes encode for regulators of the mitochondrial retrograde (RTG) pathway, which is activated in response to various physiological stresses including glutamate depletion [13], loss of respiration capacity [14], and ER stress [15]. RTG activation facilitates the transcription of several RTG-response genes that function upstream of alpha-ketoglutarate (αKG) production, including *CIT1* and *CIT2* (mitochondrial and peroxisomal citrate synthase, respectively), *DLD3* (2-hydroxyglutarate transhydrogenase), *ACO1* (mitochondrial aconitase), *IDH1* and *IDH2* (mitochondrial isocitrate dehydrogenase) [13, 14]. In WT respiratory-competent strains, some of these genes (*CIT1, ACO1, IDH1, IDH2*) are also under HAP (Heme Activator Protein) control (Fendt and Sauer, 2010). The HAP complex is a heme-activated, glucose-repressed transcriptional activator of respiratory gene expression [16]. We previously reported that canavanine alters expression of RTG-target genes in both WT and petite cells [11] both backgrounds show canavanine-induced reduction in the prototypical RTG-target genes *CIT2* and *DLD3*; *CIT1, ACO1, IDH1,* and *IDH2* expression are increased in WT, but decreased in the petite strain.

Since we see specific sensitivities in both respiratory (*rtgΔ* mutants) and non-respiratory (petite) strains with altered amino acid metabolism, we propose that the observed canavanine sensitivity is a specific type of stress that, for the purpose of this paper, we refer to simply as amino acid stress. This stress stems from the need to increase production of the absent amino acid (arginine) in order to meet metabolic requirements for protein translation, which is enhanced by even sublethal doses of canavanine since some protein molecules will become misfolded and should be replaced. Genetic backgrounds that lack robust arginine biosynthesis capabilities, like RTG mutants and petites, show reduced growth when media lacks arginine, so adding the toxic analog canavanine compounds that stress [11].

Yet the question remains: does the stress imposed by sublethal doses of canavanine inhibit growth due to the inability to generate enough arginine *per se* (which could be the case if arginine is a limiting metabolite in growth, like in petites [17]), or is sublethal toxicity due to effects on the proteome, like what occurs at much higher concentrations? Thinking more broadly - does protein misfolding affect amino acid biosynthesis? If so, then inhibiting molecular chaperones could sensitize cells to toxic amino acid exposures, offering potential value to the combination of an HSP-inhibitor and low-dose toxic amino acid analog in cancer therapeutics or as antifungal agents.

Deprivation of multiple amino acids, like in starvation conditions, causes a global suppression of protein synthesis while the expression of various stress-response genes is increased [18].

Several lines of evidence support the notion that shortage in a specific amino acid can trigger proteotoxic stress. In mammalian cells, lysine, glutamine, and leucine starvation correspond to a reduction in *HSF1* (Heat Shock transcription Factor) as well as a reduction in several heat shock protein (HSP) mRNAs. In the case of leucine, this decreased chaperone capacity sensitizes cells to proteotoxic stress [19]. However, it is not clear that *HSF1* is a regulatory factor in the amino acid starvation response. Furthermore, the cellular response to mistranslation is affected by which amino acid is being mistranslated - for example, yeast cells carrying a tRNA that substitutes serine for arginine results in a heat shock response, but proline to alanine substitution results in a minimal heat shock response [20]. Thus, the amino acid stress imposed by arginine deprivation and canavanine may be a specific response due to the inherent properties of arginine rather than an example of a more general toxic amino acid analog response.

Here we aim to explore synthetic lethality between toxic amino acids, mutations in RTG-target genes, and mutations in molecular chaperons. To investigate the connection between protein misfolding and the cellular response to depletion of either arginine or lysine, which both require αKG and glutamate for their biosynthesis, we examined the expression of RTG-target genes in mutants of heat shock proteins (HSP). We also constructed double deletion mutants - strains that lack both an RTG-target gene (*CIT1/2, DLD3, IDH1/2*) and a HSP. HSPs are chaperones that help maintain proteostasis; their expression can increase in response to a variety of stresses. HSP mutants are under constitutive proteotoxic stress, even without the addition of external stressors like heat shock or drug exposure [21, 22]. We chose four backgrounds for our experiment - two strains that lack a cytoplasmic HSP (*cpr7Δ, sse2Δ*), and two strains that lack a mitochondrial HSP (*hsp78Δ*, *mdj1Δ*). Cpr7 forms a complex with Hsp90 and has strong chaperone abilities, possibly due to its interaction with other chaperones [23] [24]. Sse2 is part of the HSP70 class of chaperone proteins, which assist in protein folding [25]. Hsp78, also part of the HSP70 class, is a mitochondrial heat shock protein that is especially important in preventing the aggregation of misfolded proteins [26]. Mdj1 is part of the HSP40 family and is a cochaperone to Ssc1, where it assists in folding, refolding, and preventing aggregation of unfolded proteins in the mitochondrial matrix. Like Hsp78, Mdj1 is also located within the mitochondria; null mutants are temperature-sensitive petites that have lost the mitochondrial genome. Mitochondrial protein import in *mdj1Δ* mutants is fine, but they are impaired in the folding of newly imported proteins [21].

Our results show that arginine deprivation and lysine deprivation have largely opposite effects on the expression of RTG-target genes in HSP mutants. RTG-target gene expression in HSP mutants under arginine or lysine deprivation has only a limited prediction value for sensitivity to toxic amino acids. Synthetic lethality can be obtained between some, but not all HSP mutants and the toxic analogs when heat shock is applied. Deletion of some of the RTG-target genes, particularly *IDH1* and *IDH2*, results in severe growth defects under sublethal analog exposure, which is exacerbated by an additional mutation in certain HSPs. In WT, canavanine and thialysine have similar effects on RTG-target gene expression; in HSP mutants, canavanine reliably reduces expression of RTG-target genes that matches the previously reported petite expression profile [11], but thialysine has a differential effect on each HSP mutant that is not a general inhibitory effect like canavanine.

## Materials and Methods

### Yeast strains

We use the WT laboratory strain BY4741. *hsp78Δ*, *sse2Δ*, *cpr7Δ, mdj1Δ, cit1Δ, cit2Δ, dld3Δ, idh1Δ,* and *idh2Δ* were of the library constructed by Giaever et al. [27]. Double knockouts were constructed by transforming a G418-resistant HSP mutant with a PCR product containing a *URA3* gene to replace either *CIT1*, *CIT2*, *DLD3*, *IDH1*, or *IDH2* and selection done on -URA, G418, and YPG media (to screen for respiratory status) followed by PCR confirmation. Primers used for transformation and confirmation can be found in Supp. Table 1. A minimum of two independent transformants with matching phenotypes in growth experiments were verified for each double mutant in order to reduce the chances of observing a phenotype due to a random additional mutation.

### Media

All experiments with canavanine were carried out in complete synthetic media (CSM) without arginine (-ARG). Similarly, experiments using thialysine were carried out on media lacking lysine (-LYS). YPD solid media contains 2% Bacto Agar, 2% Bacto Peptone, 1% Bacto Yeast Extract (Difco, Sparks, MD, USA), and 2% anhydrous dextrose (Avantor Performance Materials, Center Valley, PA, USA) dissolved in DDW. For YPD + G418 plates, G418 was added 1 ml for 400 ml. All other media (complete, YPG, media lacking uracil, or media lacking a specific amino acid with or without a toxic amino acid analog) contain 2% Bacto Agar, 0.67% Yeast Nitrogen Base (YNB) without Amino Acids (Difco Laboratories), 0.082% Complete Supplement Mixture (no amino acid drop-out) or 0.074% Complete Supplement Mixture Drop-out: ARG, LYS, or URA (Formedium). All media (excluding YPD) are brought to pH 5.8 using NaOH pearls (Bio-Lab LTD, Jerusalem, IL, USA), autoclaved, and 2% glucose is added after autoclaving. Liquid media were prepared in the same fashion as solid media with the exclusion of Bacto Agar.

### RNA purification and RT-qPCR measurement of mRNA

For each strain (BY4741, *cpr7Δ*, *sse2Δ*, *hsp78Δ*, and *mdj1Δ*), single colonies were picked from YPD agar plates and patched onto -ARG, -ARG + 1 μg/ml canavanine, -LYS, or -LYS + 6 μg/ml thialysine solid media and grown for four days in 30°C, for a total of three independent isolates for each strain in each condition. Cells were scraped from solid media, flash frozen with liquid nitrogen, and stored in -80°C if not immediately harvested. Total RNA was isolated using the RNeasy kit (Qiagen) and the MasterPureTM Yeast RNA Purification Kit (Epicentre). cDNAs were synthesized with the FastQuant RT Kit (with gDNase) according to the manufacturer’s protocol. Real-time PCRs were done in a 10 μl volume using the Absolute quantitative PCR SYBR Green mix (Sigma) in a 96-well plate. Two technical repeats of each sample were used, with three independent isolates for each genotype. The real-time PCR primers used: forward 5′–3′ primer for *ALG9*: TCACAAGAGCATGCTTAGGC; reverse 5′–3′ primer for *ALG9*: AAGCTGCCTGCAATTTCACG; forward 5′–3′ primer for *CIT2*:GGTTCCAACTCAAGCGCAAG; reverse 5′–3′ primer for *CIT2*: ACCCGTTCAAACCTGATGCA; forward 5′–3′ primer for *DLD3*: ACGTCAGGGTCCAATAAGAGACAC; reverse 5′–3′ primer for *DLD3*: CAAACCGGCTGCGTTTAATCTCTC; forward 5′–3′ primer for *ACO1*: AGTAACTGCGTTCGCCATTG; reverse 5′–3′ primer for *ACO1*: AGGCAAACCATCACCATGTG; forward 5′–3′ primer for *IDH1*: TTGTCGACAATGCCTCCATG; reverse 5′–3′ primer for *IDH1*: TCAAAGCAGCGCCAATGTTG; forward 5′–3′ primer for *IDH2*: TTGCCGGTCAAGATAAAGCG; reverse 5′–3′ primer for *IDH2*: TGTTTTCTGGACCTGATGCG; forward 5′–3′ primer for *CIT1*: TGGTCGTGCCAATCAAGAAG; reverse 5′–3′ primer for *CIT1*: AAAACCGCATGGCCATAACC. *ALG9* was used as the internal control with RNA levels of the genes of interest normalized to *ALG9* levels [28]. PCRs were initiated at 95°C for 20 s in the holding stage. The cycling stage consisted of 3 s at 95°C, 30 s at 60°C, and 15 s at 95°C, repeated for 40 cycles. Melt curve stage consisted of 15 s at 95°C followed by 1 min at 60°C and finally 15 s at 95°C. Fold-changes in mRNA expression of RTG-target genes and standard deviation were calculated using the Δ/Δ Ct method. For HSP mutants grown on -ARG, fold-changes are relative to WT expression on -ARG. For HSP mutants grown on -LYS, fold- changes are relative to WT expression on -LYS. WT expression is a “self” comparison reflecting expression levels under canavanine exposure (WT CAN versus WT -ARG), or expression levels under thialysine exposure (WT AEC versus WT -LYS). Similarly, HSP mutant expression under canavanine and thialysine exposure is also a “self” comparison that compares expression under analog exposure to expression on media without drug. Heat map visualization of fold-change in each RTG-response gene and subsequent cluster analysis was done with Morpheus, https://software.broadinstitute.org/morpheus; cluster analysis was constructed by calculation of Euclidean distance based on complete linkage for each RTG-target gene of a strain/condition.

### Growth assays

For spot assays on solid media, using a 96-well tissue culture plate (Jet Biofil), a single colony undergoes a tenfold serial dilution. Using a 6-by-8-column metal prong, approximately 1 μl from each well is pronged onto desired media. After allowing suspended cells to fully absorb into the plate, the plate is put into 30°C or 37°C and growth photographed at the indicated time points. For growth assays in liquid media, a single colony was placed into a 96-well tissue culture plate containing 200 µl YPD in each well and grown overnight in 30°C at 200 RPM. The following day, after vigorous shaking to ensure suspension of cells, 5 µl from each culture was diluted into 100 µl DDW in a new 96-well plate. From this “water culture”, 3 µl of culture was transferred to 200 µl of either -ARG or -LYS. Using a TECAN plate reader, OD_600_ was measured every hour with agitation. Growth curves depict the average of at least two independent cultures for each genotype; the OD at each time point represents the average of nine technical repeats for that well (OD is read at nine different locations within a well) (Supp. Table 2).

## Results

### Effect of toxic amino acid analogs on RTG-response gene expression in WT

To test which RTG-target genes are affected at the transcript level by exposure to toxic amino acid analogs, WT cells were grown on solid media containing a sublethal concentration of canavanine (1 µg/ml) (WT CAN) and thialysine (6 µg/ml) (WT AEC). *CIT2* and *DLD3* are reduced in response to both toxic amino acid analogs. The expression of *CIT1* is maintained under canavanine exposure, and *ACO1*, *IDH1,* and *IDH2* expression are induced by canavanine (Fig. 1A); *IDH1* and *IDH2* show particularly strong increases in expression (2.7-fold and 16-fold, respectively). The same WT strain has a slightly different response pattern to thialysine - *CIT1* and *ACO1* expression are modestly reduced, but similar to canavanine exposure, *IDH1* and *IDH2* show increased expression (1.3-fold and 4.8-fold, respectively) (Fig. 1B).

**Fig. 1.**
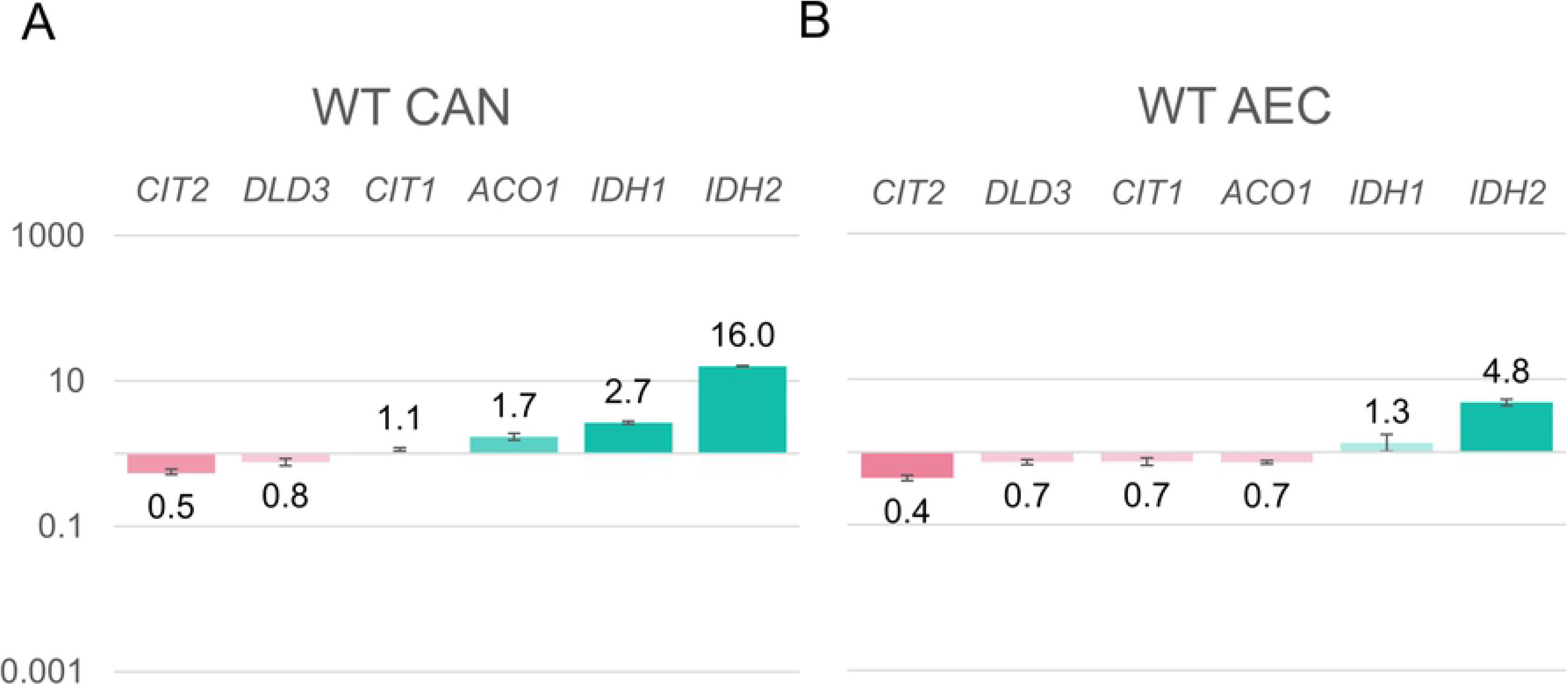
Canavanine and thialysine alter expression of RTG-target genes in WT. RNA was extracted from strain grown on solid media with or w/o exposure to toxic amino acids. Fold change was calculated using the Δ/ΔCt method with *ALG9* as a reference gene. A) Bar chart of WT RTG-response gene mRNA expression grown on solid glucose –arginine + 1 µg/ml canavanine (WT CAN) relative to WT grown on glucose –arginine. B) Bar chart of WT RTG- response gene mRNA expression grown on solid glucose –lysine + 6 µg/ml thialysine (WT AEC) relative to WT grown on glucose –lysine. Three biological replicates were analyzed for each condition.

### Arginine deprivation and lysine deprivation have opposite effects on RTG-target gene expression in HSP mutants

Next, we tested the expression of the same RTG-target genes in strains with inherent misfolding stress under arginine deprivation. This was done in strains mutated in HSP mutants that are missing a chaperone protein. When grown on -ARG, relative to WT, all RTG-target genes are induced in *hsp78Δ* and *sse2Δ* (Fig. 2A). With the exception of *CIT2*, all RTG-target genes are induced in *mdj1Δ.* Although the reduction in *CIT2* is relatively small, being a petite strain, we expected *CIT2* to be higher in this background relative to WT; nonetheless, to the best of our knowledge, we are the first to measure *CIT2* expression in this background and under these conditions. *cpr7Δ* shows a different RTG-response gene expression profile than the other three HSP mutants; it has significantly reduced *CIT2*, *DLD3,* and *ACO1; CIT1* expression is equal to WT. Of all measured RTG-response genes, only *IDH1* and *IDH2*, which are strongly induced in *mdj1Δ, hsp78Δ*, and *sse2Δ,* show increased expression in *cpr7Δ* (Fig. 2A).

**Fig. 2.**
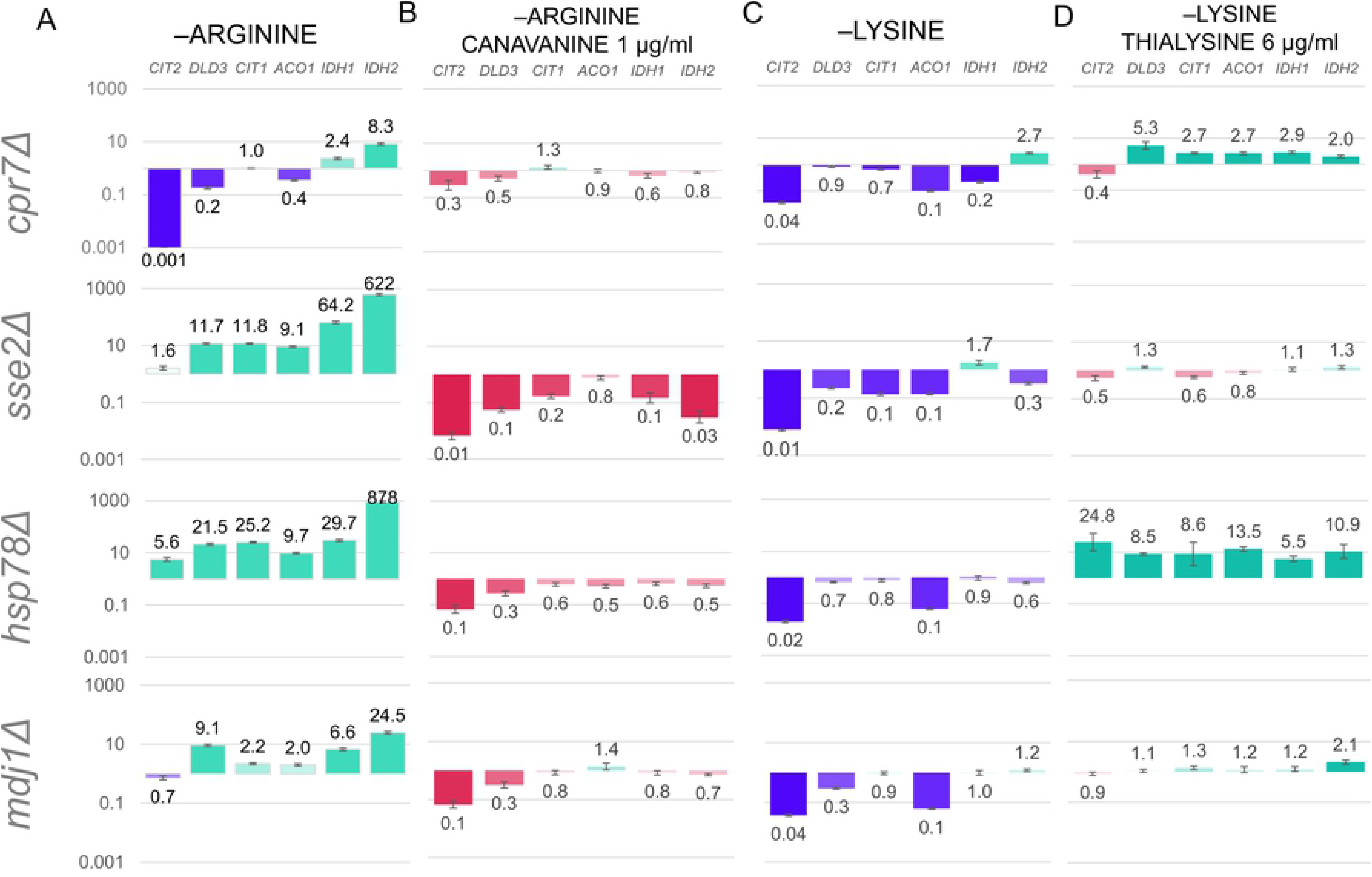
RTG-target gene expression in HSP mutants grown under arginine deprivation +/- canavanine exposure, and lysine deprivation +/- thialysine exposure. RNA was extracted from the indicated strains grown on solid media. Fold change was calculated using the Δ/ΔCt method with *ALG9* as a reference gene. RTG-gene expression of HSP mutants grown on glucose – arginine is relative to WT grown on glucose –arginine. A) Bar chart shows fold change of measured RTG-target genes under arginine deprivation. B) Bar chart shows fold change of measured RTG-target genes under arginine deprivation + 1 µg/ml canavanine. C) Bar chart shows fold change of measured RTG-target genes under lysine deprivation. D) Bar chart shows fold change of measured RTG-target genes under lysine deprivation + 6 µg/ml thialysine.

Since lysine and arginine both require αKG and glutamate for their biosynthesis, we measured RTG-target gene expression in mutants grown without lysine; surprisingly, lysine deprivation has almost the opposite effect of what we see in the same mutants grown without arginine. All HSP mutants show a considerable reduction in *CIT2* and *ACO1,* with variable levels of reduction in *DLD3* and *CIT1* (Fig. 2C). *hsp78Δ* also shows reduced *IDH1* and *IDH2*, although when taking into account the standard deviation, *IDH1* expression may be equal to WT under the same conditions. Surprisingly, *mdj1Δ* maintains the same pattern as the respiratory-proficient mutants and also shows reduced RTG-target gene expression relative to WT. The only RTG- target genes that show increased expression under lysine deprivation are the two mitochondrial isocitrate dehydrogenase genes: *IDH1* in *sse2Δ* (1.7-fold), and *IDH2* in *cpr7Δ* (2.7- fold) and *mdj1Δ* (1.2) (Fig. 2C).

To get a more comprehensive view of the effect of mild protein misfolding on the expression of RTG-target genes we applied a clustering algorithm (Euclidean distance based on complete linkages) to the fold-changes (Figs. 1, 2A and 2C) of RTG-target gene expression of HSPs -ARG, HSPs -LYS, WT CAN, and WT AEC. *hsp78Δ* and *sse2Δ* grown on -ARG form their own cluster due to their robust expression of all measured RTG-target genes. The remaining strains/conditions are part of the second cluster. *cpr7Δ* -ARG immediately clusters with WT CAN. All HSP mutants grown on -LYS, including *mdj1Δ,* cluster with WT AEC; *cpr7Δ* -LYS again clusters immediately with WT AEC (Fig. 3). *mdj1Δ* -ARG is within this large cluster, but separates out from the other strains/conditions, a reflection of its inherent arginine biosynthesis issues.

**Fig. 3.**
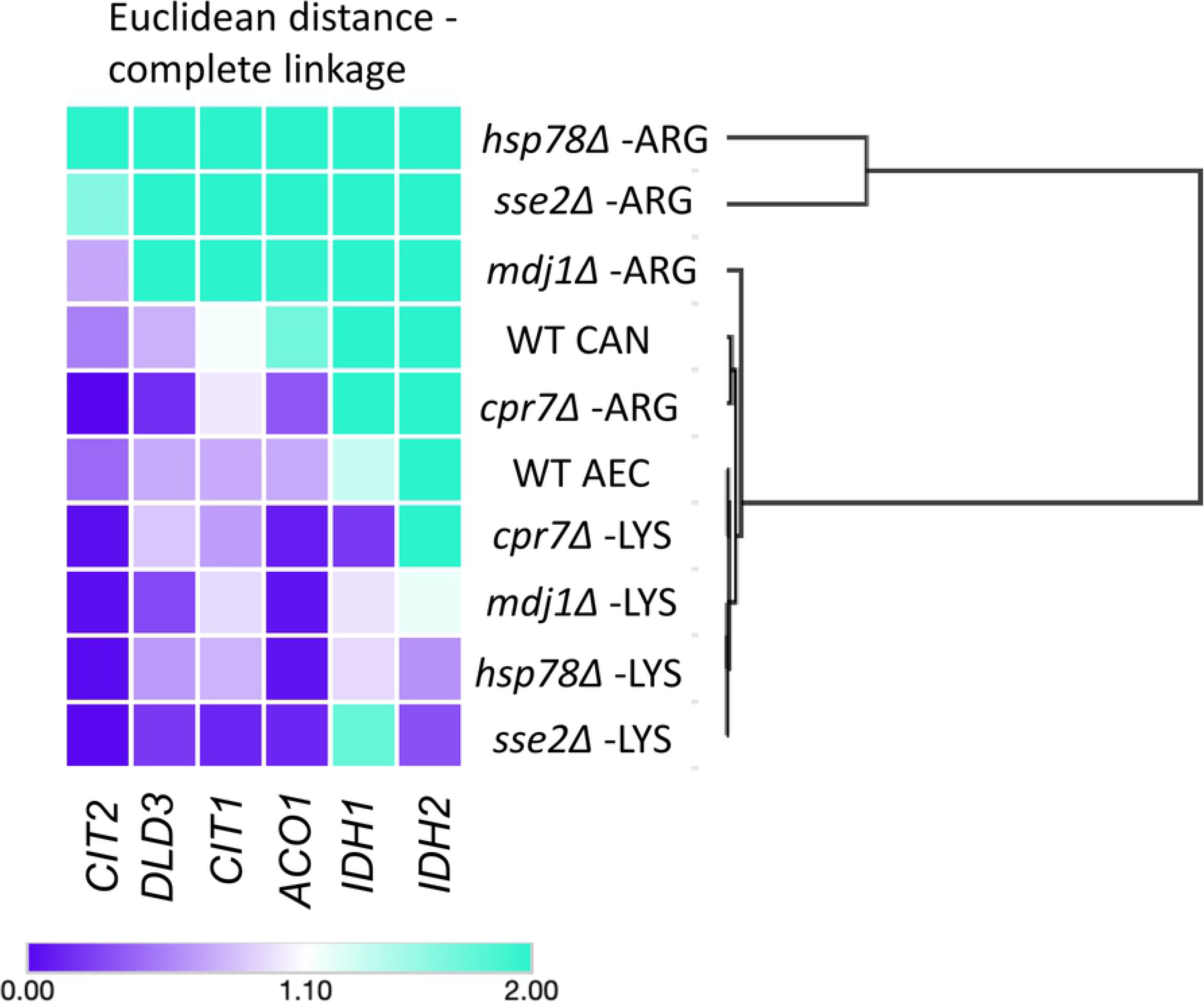
Hierarchical clustering of WT under toxic analog exposure and HSP mutants under arginine deprivation and lysine deprivation. Heat map depicting fold-change of each measured RTG-target gene (from Figs. 1, 2A, and 2C) with applied hierarchical Euclidean clustering based on complete linkage.

### Effect of toxic amino acid analogs on RTG-response gene expression in HSP mutants

Respiratory-deficient yeast (petites) have RTG-dependent increases in the expression of RTG- target genes in comparison with the respiratory WT. We previously showed that low-dose canavanine exposure leads to reduced expression of all measured RTG-target genes in a petite strain [11]. The above expression data demonstrates that HSP mutants show altered expression of RTG-target genes relative to WT, so we wanted to know how canavanine and thialysine exposure change those expression patterns. Low-dose canavanine exposure has a reliable effect in reducing RTG-target gene expression; *CIT2* and *DLD3* are reduced in all HSP mutants (Fig. 2B) and WT (Fig. 1). HSP mutants show RTG-target gene expression that resembles that of the petite rather than WT [11]. *IDH1* and *IDH2* also show reduced expression in all HSP mutants (Fig. 2B).

In WT, the expression pattern induced by canavanine and thialysine is quite similar (Fig. 1), and in HSP mutants, canavanine exposure has a reliable effect in suppressing RTG-target gene expression (Fig. 2B); however, thialysine does not have a consistent effect on RTG-target gene expression in HSP mutants (Fig. 2D). *hsp78Δ* shows a robust induction of all RTG-target genes under thialysine exposure - the opposite of its response to canavanine. Thialysine exposure reduces *CIT2* expression in *cpr7Δ*, yet all other RTG-target genes show two-fold and greater induction. Of the four growth conditions for mRNA expression analysis, RTG-target gene expression in *sse2Δ* shows the least response to thialysine, with *CIT2* showing the greatest reduction (0.5-fold reduction compared to growth in -LYS). *mdj1Δ* also shows a mild response to thialysine; there is a slight reduction in *CIT2* (0.9-fold decrease), but the expression of all other RTG-target genes is maintained or slightly increased despite the presence of the toxic lysine analog, and *IDH2* shows the greatest induction (2.1-fold increase).

When taking into account the effect of an HSP mutation on the expression of RTG-target genes under either arginine or lysine deprivation (Figs. 2A and 2C), and then the effect of toxic amino acid exposure in these backgrounds (Figs. 2B and 2D), we often observe reverse relationships. For example, compared to WT, both *hsp78Δ* and *sse2Δ* show robust RTG-target gene expression under arginine deprivation, but canavanine exposure causes a considerable reduction in all measured genes. Likewise, *hsp78Δ* has reduced expression of RTG-target genes under lysine deprivation, but the same genes are induced by thialysine exposure. *cpr7Δ* also shows a similar trend under lysine deprivation (reduced RTG-target gene expression) and thialysine exposure (increased RTG-target gene expression).

### Functional connection between amino acid stress and proteotoxic stress

We found that in WT, both canavanine and thialysine exposure alter the expression of RTG- target genes in a similar manner under our conditions. Our prediction was that HSP mutant genotypes in which RTG-target gene expression under arginine or lysine deprivation was similar to WT CAN or WT AEC would show increased sensitivity to the respective toxic analog.

However, we found that both analogs have a different effect on RTG-target gene expression in HSP mutants than in WT (Figs. 1, 2B and 2D). In addition, the expression of RTG-target genes is just one possible determinant of toxic amino acid tolerance; we have previously shown that respiration status also determines canavanine tolerance, but not thialysine tolerance [11]. We directly tested the sensitivity of HSP mutants on sublethal doses of canavanine and thialysine. Due to the petite nature of *mdj1Δ*, we expect it to be highly sensitive to 1 µg/ml canavanine regardless of its RTG-target gene expression in -ARG, and it is (Fig. 4A). Although we do not detect growth differences between WT and respiratory HSP mutants on solid 1 µg/ml canavanine (the concentration used for mRNA expression analysis), increasing the concentration to 2 µg/ml reveals a clear growth inhibition in *cpr7Δ* that is not present in either *hsp78Δ* or *sse2Δ* (Fig. 4A). Thus, respiratory mutants that show elevated expression of all RTG- target genes under arginine deprivation (*hsp78Δ*, *sse2Δ*) are less sensitive to canavanine than *cpr7Δ*. Respiratory HSP mutants show no growth defects from heat shock when grown without arginine. When proteotoxic stress is increased by combining 2 µg/ml canavanine with heat shock, both *hsp78Δ* and *cpr7Δ* show a clear growth defect (Fig. 4C).

**Fig. 4.**
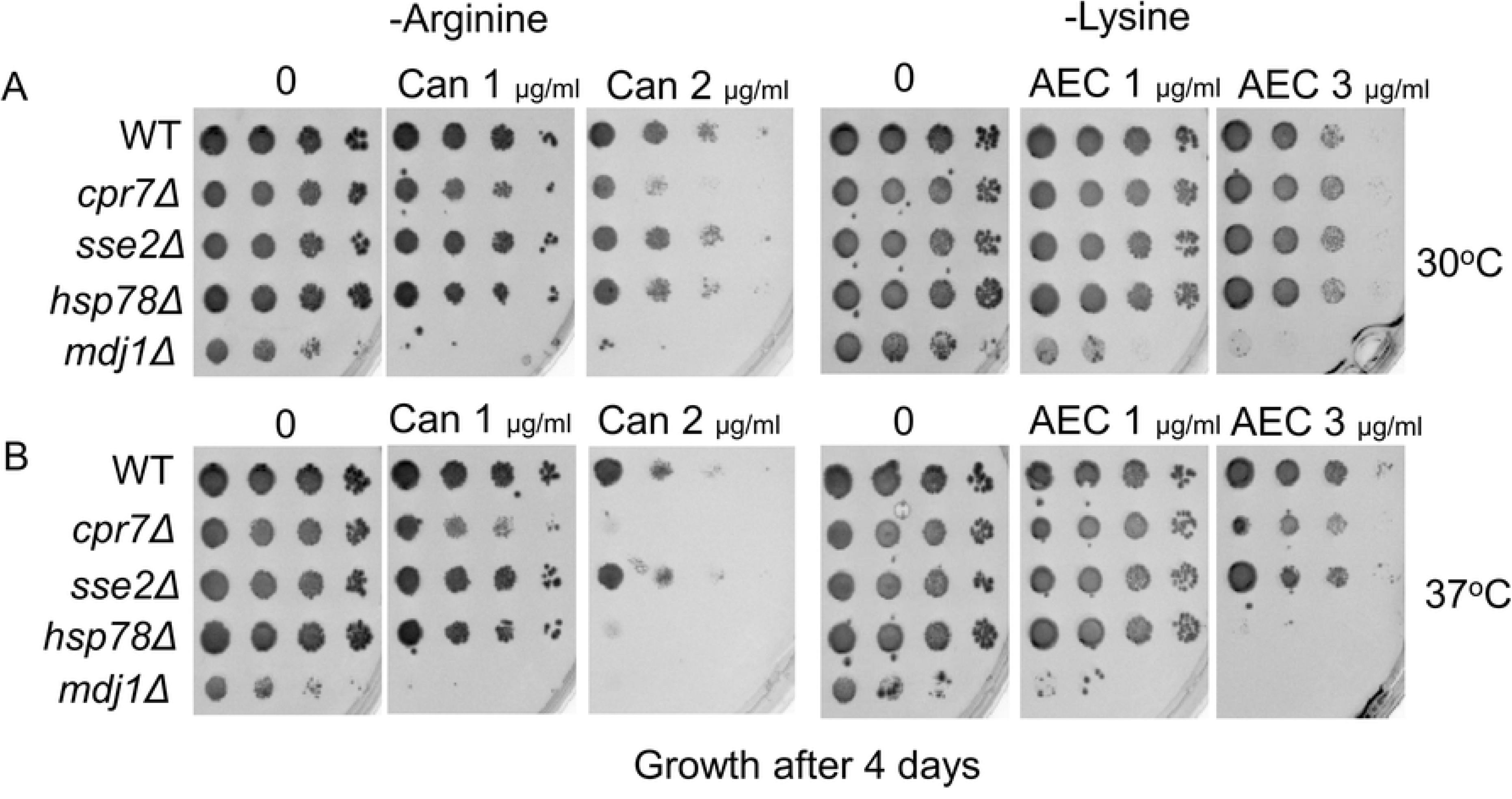
Under heat shock conditions*, cpr7Δ* and *hsp78Δ* are sensitive to canavanine and thialysine. A single colony was placed in 100 µl DDW and 10-fold dilutions performed, followed by transfer of approximately 1 µl of suspended cells to plates using a 6X8 metal pronger, a procedure we call spot assay. Here the spot assay shows growth of WT and HSP mutants on A) glucose –arginine +/ – canavanine at 30°C; B) glucose –lysine +/ – thialysine at 30°C; C) glucose –arginine +/ – canavanine at 37°C; and D) glucose –lysine +/ – thialysine at 37°C. Growth after four days incubation.

Under lysine deprivation conditions, yeasts, including petites, can tolerate higher concentrations of thialysine. All respiratory HSP mutants grow as well as WT (Fig. 4B) on 3 µg/ml thialysine at 30°C. As a petite, *mdj1Δ* grows slower than the respiratory strains, so it shows a lag in growth after only four days of incubation; after a week, *mdj1Δ* grows to WT level (Supp. Fig. 1), and can tolerate as much thialysine as WT. At 6 µg/ml thialysine (the concentration used for mRNA expression analysis), *hsp78Δ* and WT show a comparable, albeit small, reduction in growth; surprisingly, deletion of *CPR7* and *SSE2* renders a slight improvement in the ability to form colonies at this concentration (Supp. Fig. 1). Under the combined stress of heat shock and 3 µg/ml thialysine, *hsp78Δ* fails to grow, and *cpr7Δ* also shows increased sensitivity; however, heat shock does not diminish growth in either WT or *sse2Δ* at this concentration (Fig. 4D). *mdj1Δ* is a temperature-sensitive mutant even under ideal nutritional conditions (YPD).

In summary, only *sse2Δ* shows WT-level growth under all tested conditions; *hsp78Δ* is sensitive to both thialysine and canavanine, but only when combined with the additional stress of heat shock; *cpr7Δ* is the most canavanine sensitive of the respiratory strains, but in the absence of heat shock, tolerates thialysine better than the WT. *mdj1Δ* is highly and specifically sensitive to canavanine - given sufficient incubation time, petites can tolerate thialysine as well as WT.

These results highlight the complex interactions between single amino acid deprivation, low-dose toxic analog exposure, mitochondrial status, and genotype.

### Dissection of RTG-response genes in HSP mutants

Finally, we created a set of double mutants in which we systematically knocked out each RTG- target gene in the HSP mutant backgrounds, except for *ACO1*, as we were unable to delete this gene using our transformation method. We compared growth of HSP/RTG-target gene double mutants to their respective single mutants (*cpr7Δ*, sse2*Δ, hsp78Δ, mdj1Δ, cit1Δ*, *cit2Δ, dld3Δ, idh1Δ,* and *idh2Δ*). Of the RTG-target gene single mutants, only *idh1Δ* and *idh2Δ* show sensitivity to canavanine and thialysine (Supp. Fig. 2). This corresponds to deficiencies in arginine and lysine production evident by their reduced growth on -ARG and -LYS (Supp. Fig. 3). Growth on canavanine reveals a possible negative genetic interaction (the double mutant grows worse than each of the single mutants) between *idh1Δ/idh2Δ* and *hsp78Δ* and *sse2Δ* when combined with heat shock (Fig. 5, Supp. Fig. 2). *hsp78Δidh2Δ* and *mdj1Δidh2Δ* also show increased sensitivity under thialysine exposure relative to the single mutants (Supp. Fig. 2).

**Fig. 5.**
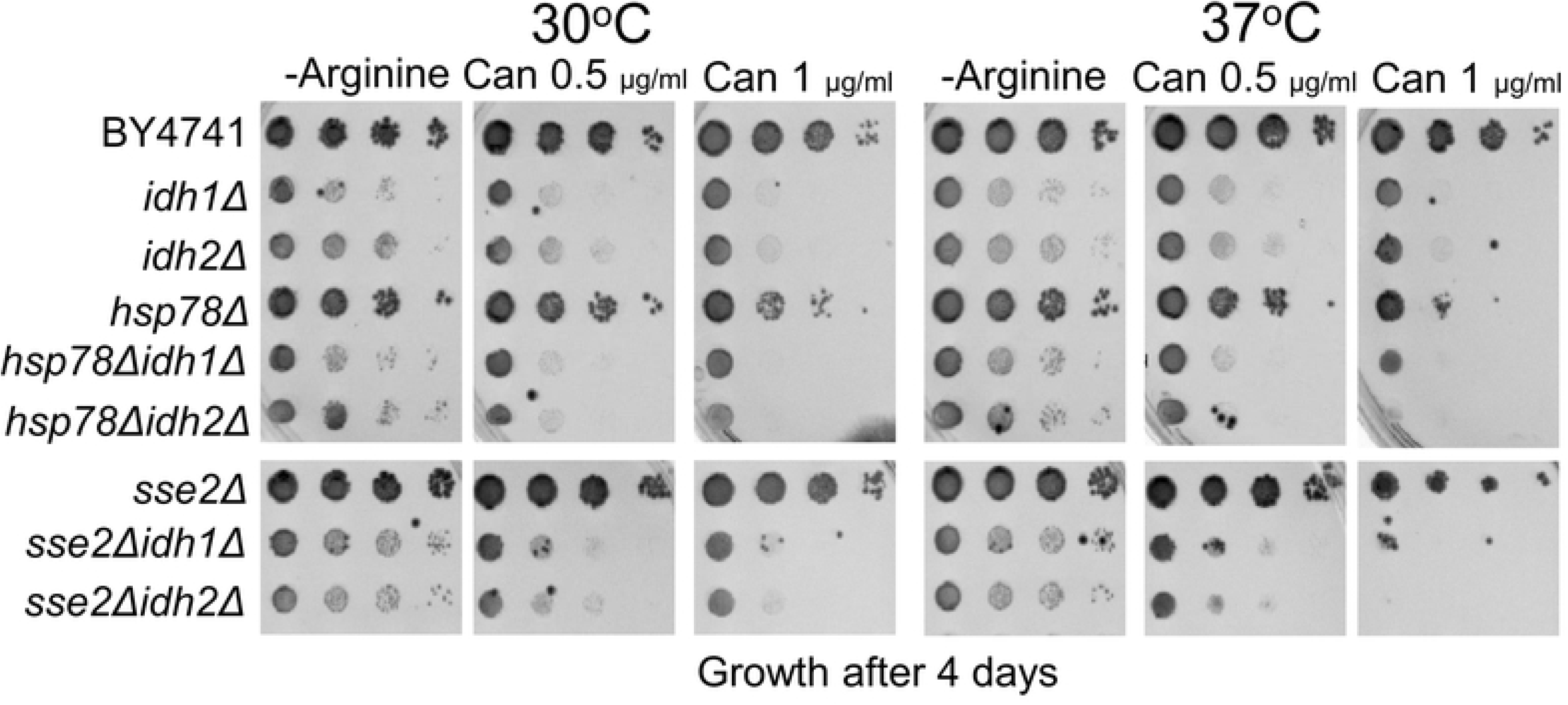
*IDH1* and *IDH2* are required for canavanine tolerance under heat shock conditions in the HSP mutants *sse2Δ* and *hsp78Δ*. A spot assay was done as described in the legend of Fig.4. Spot assay showing growth of WT, *idh1Δ*, *idh2Δ*, the HSP single mutants *sse2Δ* and *hsp78Δ*, and their respective *idh1Δ/2Δ* double mutants on glucose –arginine +/ – canavanine after four days incubation in 30°C (left panel) or 37°C (right panel).

So far, we demonstrated genetic interactions between chaperones and *IDH1*, *IDH2*, and *CIT1*, which function in the TCA cycle, as well as mitochondrial chaperone mutants and *CIT2,* the prototypical RTG-response gene that is located in the peroxisome and part of the glyoxylate cycle. *DLD3,* which converts the oncometabolite D2HG to αKG, is not part of the TCA cycle but is also strictly under RTG control like *CIT2*. In solid media at 30°C, *HSP78* also reveals a negative genetic interaction with *DLD3* on as little as 0.5 µg/ml thialysine; upon heat shock, this synthetic lethality is apparent on both canavanine and thialysine (Supp. Fig. 2). To the best of our knowledge this is the first report on genetic interactions between HSP proteins and RTG- target genes.

### Effect of arginine deprivation and lysine deprivation on growth dynamics of HSP/RTG-target gene mutants

Growth dynamics can be assessed at a higher resolution by following OD in liquid media, so we followed the growth of WT and HSP single mutants for 48 hours in -ARG and -LYS at 30°C. Notably, WT has a slightly elongated lag phase in -ARG compared to -LYS (Supp. Fig. 4); increased time in lag phase is characteristic of metabolic remodeling and is often observed in the switch from rich to minimal media, with different genotypes requiring more or less time. *mdj1Δ* spends the longest time in lag phase and shows more growth inhibition by arginine deprivation compared to lysine deprivation (Supp. Fig. 4).

Next, we followed the growth of respiratory *HSP/IDH1/IDH2* double mutants for 48 hours in - ARG and -LYS at 30°C. Notably, *idh1Δ* and *idh2Δ* show greater growth inhibition compared to WT when grown in -LYS as opposed to growth in -ARG, although both conditions reveal obvious growth defects (Fig. 6). Deletion of *IDH1* has the greatest negative effect on growth compared to WT in both -ARG and -LYS, and it has still not reached stationary phase after 48 hours growth in -LYS (Fig. 6, Supp. Fig. 4). All three respiratory *HSP/IDH2* double mutants show severe growth defects under lysine deprivation; *cpr7Δidh2Δ, sse2Δidh2Δ,* and *hsp78Δidh2Δ* show even worse growth than *idh2Δ* or the parent single mutant alone (Fig. 6, Supp. Fig. 5).

**Fig. 6.**
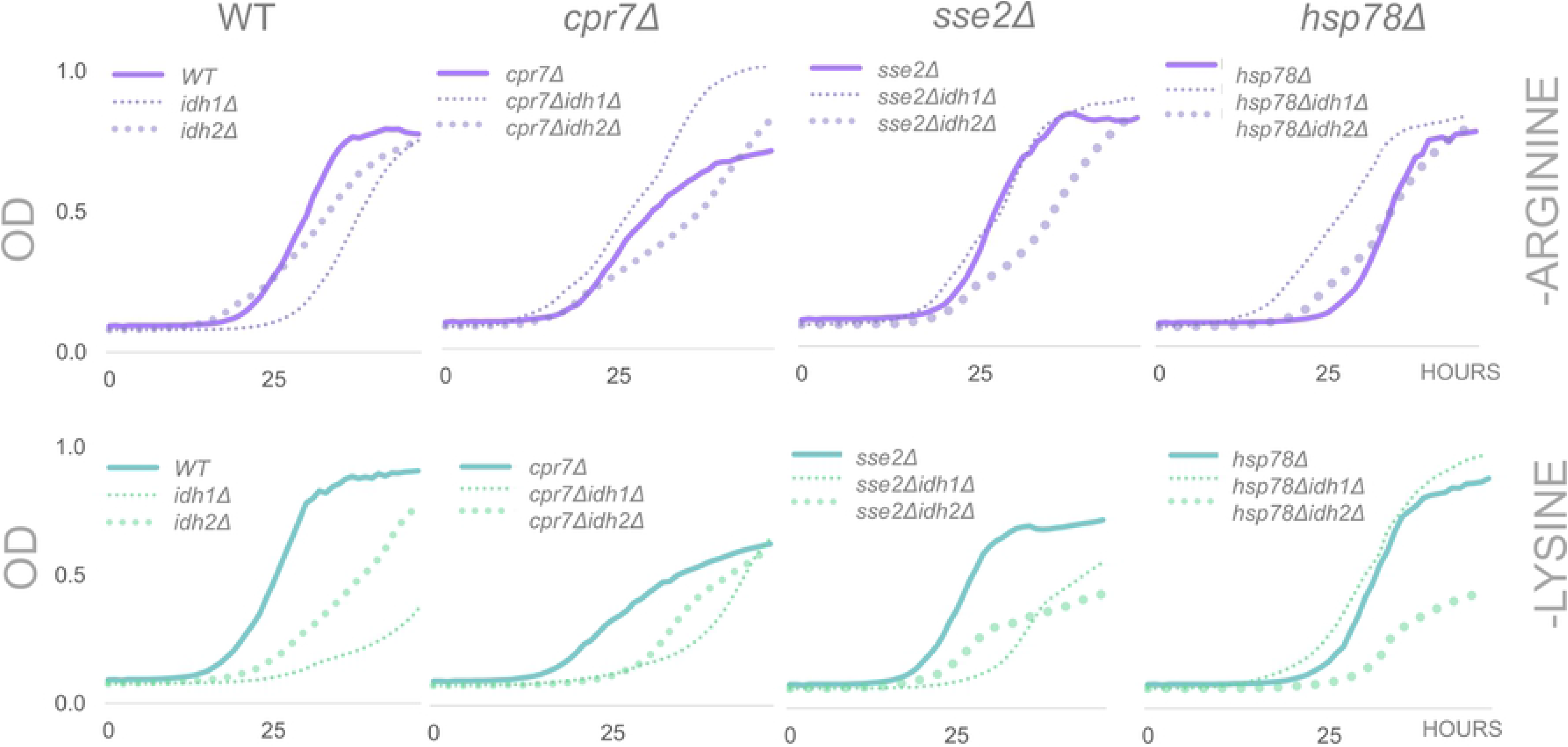
Growth curves of *IDH1/IDH2* deletion in respiratory backgrounds. WT, *idh1Δ*, *idh2Δ*, single HSP mutants (*cpr7Δ*, *sse2Δ,* and *hsp78Δ*) and their respective *idh1Δ/2Δ* double mutants grown in glucose –arginine (purple) and glucose –lysine (green) for 48 hours. OD was read every hour. OD measurements are found in Supp. Table 2.

The spot assay does not detect canavanine/thialysine-induced growth defects from *CIT1* and *CIT2* deletion in WT. However, growth on 1 µg/ml thialysine suggests a potential negative genetic interaction between HSP mutants and *CIT1*/*CIT2* (Supp. Fig. 2). We followed the growth of *HSP/CIT1/CIT2* double mutants. Under arginine deprivation, deletion of *CIT1* does not negatively affect growth rate or final cell count at stationary phase in either WT or HSP mutants (Supp. Fig. 6); in the WT, *CIT2* deletion decreases final cell count at stationary phase; with the exception of *mdj1Δ*, this effect is not compounded by the additional deletion of an HSP gene - *cit2Δ*, *cpr7Δcit2Δ*, *sse2Δcit2Δ*, and *hsp78Δcit2Δ* all show similar growth curves under arginine deprivation (Supp. Fig. 6), and grow as well or better than the single HSP mutant parent.

Lysine deprivation leads to a greater reduction in final cell count in both *cit1Δ* and *cit2Δ* compared to WT (Supp. Fig. 6). In *cit1Δ*, this effect is not compounded by the additional deletion of an HSP gene - *cit1Δ*, *cpr7Δcit1Δ*, *sse2Δcit1Δ*, *hsp78Δcit1Δ* and *mdj1Δcit1Δ* all show similar growth curves in liquid media lacking lysine (Supp. Fig. 6).

Under arginine deprivation, all *hsp78Δ* double mutants (*hsp78Δcit1Δ, hsp78Δcit2Δ, hsp78Δidh1Δ, hsp78Δidh2Δ)* grow as well or better than the HSP single mutant alone (Fig. 6, Supp. Fig. 6). Yet, under lysine deprivation, three of the four double mutants grow worse than *hsp78Δ*; only *hsp78Δidh1Δ* grows as well as the single mutant (Fig. 6, Supp. Fig. 6, Supp. Table 2). Notably, deletion of *CIT1* (mitochondrial citrate synthase) from the *mdj1Δ* background does not have a deleterious effect on growth under either arginine deprivation or lysine deprivation, but deletion of *CIT2* (peroxisomal citrate synthase) results in a severe growth defect in both - ARG and -LYS (Supp. Fig. 6).

## Discussion

Arginine deprivation therapy combined with canavanine is being explored as a current approach to specifically target and tamper with the metabolism of cancer cells [9, 10]. Canavanine-induced stress on the proteome through protein misfolding and subsequent ER stress has been documented and continues to be researched for its potential therapeutic benefits [5, 6, 8, 12]. Our previous work has demonstrated that low-dose canavanine exposure induces stress on the mitochondria related to arginine biosynthesis, and that certain genetic backgrounds and physiologic conditions are synthetically lethal when combined with extremely low doses of canavanine [11]. The work encompassed here sheds light on synthetic lethal conditions between mutated heat shock proteins, arginine/lysine deprivation, RTG-target genes, and toxic analog exposure that could possibly be exploited as therapeutic options or for their fungicidal/bactericidal ability.

To test if RTG-target gene expression links single amino acid deprivation to protein misfolding stress, we compared the mRNA signatures of WT exposed to low-dose amino acid analogs and HSP mutants grown on a complete supplement media lacking either arginine or lysine. We previously showed that both respiratory mutants and RTG-pathway mutants are extremely sensitive to canavanine [11]; here we test our hypothesis that, if the canavanine sensitivity observed at sublethal doses is a result of protein misfolding stress, then HSP mutants grown under arginine deprivation conditions will show a similar RTG-target gene expression profile to WT grown with canavanine; likewise, HSP mutants grown under lysine deprivation should show a similar RTG-response gene expression profile to WT grown with thialysine. Our data show that the majority of HSP mutants do show a similar RTG-expression profile under the specific conditions (Fig. 3); however, relative to WT, all of the mutants show a clearly different RTG- target gene expression response to the toxic analogs (Fig. 2).

Our data indicate that which HSP (Cpr7, Sse2, Hsp78, or Mdj1) and which amino acid (arginine or lysine) is lacking affect RTG-target gene expression. Applying a cluster analysis to HSP mutants grown under arginine or lysine deprivation groups six of the eight HSP cases with WT CAN and WT AEC; *hsp78Δ* and *sse2Δ* under arginine deprivation are the exceptions. Of note, our data show that *IDH1* and *IDH2* are upregulated in WT in response to stress imposed by sublethal doses of both canavanine and thialysine, as well as in all HSP mutants under arginine deprivation (Fig. 1 - 3). Thus increased *IDH1*/*IDH2* expression seems to be a feature of low-dose analog exposure, as well as arginine deprivation, which may or may not be mediated through the RTG pathway given the differential expression of *CIT2* and *DLD3* under arginine deprivation conditions (Fig.s 1, 2). However, the cluster analysis suggests that lysine deprivation - when arginine is present - induces an arguably stronger phenotype defined by reduced *CIT2*, *DLD3, CIT1,* and *ACO1* expression in all tested HSP mutants, similar to the RTG-target gene expression profile of WT AEC (Fig. 3). The heat map also reveals another pattern among the ten strains/conditions - *IDH2* is frequently upregulated, and *CIT2,* along with *DLD3*, is frequently downregulated. The simplest explanation is that both arginine deprivation and lysine deprivation are stressful conditions, but two of our tested strains, *hsp78Δ* and *sse2Δ*, are capable of full and robust activation of RTG-target genes under arginine deprivation conditions, in which case we cannot make the argument that up or down-regulation of RTG-target genes indicates protein misfolding stress, especially considering the differential responses of HSP mutants to canavanine and thialysine.

Under arginine deprivation, *cpr7Δ* has an RTG-target gene expression profile most similar to WT CAN (Fig. 1, 2A, 3). Of the respiratory mutants, *cpr7Δ* also shows the greatest sensitivity to canavanine, but only as more stress is applied (Fig. 4A, 4C). Closer analysis of the four HSP mutants used in this study shows that Cpr7, a cytoplasmic chaperone itself, interacts with many other chaperones, and of the four tested HSP mutants it has by far the most physical and genetic interactions documented on SGD (Yeastgenome.org, Supp. Fig. 7). Thus, deletion of *CPR7* likely affects protein homeostasis at multiple molecular points, which could mean that *cpr7Δ* has the highest degree of proteotoxic stress; if sublethal canavanine exposure attenuates protein homeostasis, *cpr7Δ*’s RTG-target gene expression under arginine deprivation and subsequent canavanine sensitivity support this idea. *cpr7Δ’*s inability to activate the RTG pathway under arginine deprivation (evidenced by reduced *CIT2* and *DLD3* expression) may also explain *cpr7Δ’*s canavanine sensitivity.

Lysine deprivation consistently results in downregulation of RTG-target genes in HSP mutants (Fig. 2C). Although the cluster analysis of WT AEC and HSP mutants on -LYS does not predict thialysine sensitivity the same way *cpr7Δ* clustering with WT correlates to its canavanine sensitivity (both WT and HSP mutants show basically the same sensitivity to thialysine after a week’s incubation at 30°C, Supp. Fig. 1), we do see that HSP/RTG-target gene double mutants show more severe growth defects under lysine deprivation compared to arginine deprivation (Fig. 6, Supp. Fig. 2, 5 & 6). This may indicate that the upregulation of RTG-target genes induced by arginine deprivation can compensate for deletions in a single RTG-target gene (excluding *idh1Δ*/*idh2Δ*), presumably because the rest of the target genes are sufficiently expressed. On the other hand, under conditions of lysine deprivation, RTG-target genes are downregulated, so the loss of a single gene within the pathway is more noticeable if that pathway is in fact required for growth in such conditions. It’s interesting to note that the cluster analysis groups the mitochondrial chaperone mutants *hsp78Δ* and *mdj1Δ* together under lysine deprivation conditions, yet *mdj1Δ* -ARG is separate from the other strains/conditions within this large cluster. This reflects the innate arginine biosynthesis issues of petite strains [11, 29] although petites have deficits in the synthesis of other amino acids (glutamate, glutamine, and leucine) [17], lysine is not one of them.

Of the tested RTG-target gene single mutants, only *idh1Δ* and *idh2Δ* show sensitivity to both toxic amino acid analogs, as well as growth inhibition on -ARG and -LYS (Supp. Fig. 2 & 3).

Dissection of synthetic effects between protein misfolding mutants and RTG-target gene mutants via creation of HSP/RTG-target gene double mutants reveals a potential cell-wide role for mitochondrial isocitrate dehydrogenase in the protein misfolding stress response, as both mitochondrial and cytoplasmic chaperone mutants show increased sensitivity to the combined stress of canavanine and heat shock when missing *IDH2* (Fig. 5, Supp. Fig. 2). We detect a negative synthetic genetic interaction between *HSP78* and *IDH2,* as well as *SSE2* and *IDH2* when the double mutants are exposed to the highest stress conditions on solid media (arginine deprivation + canavanine + heat shock), and possibly under -LYS + thialysine on solid media too (Supp. Fig. 2). We measured OD in liquid -ARG/-LYS, and we see clear additional growth defects in HSP mutants lacking *IDH2* when lysine is absent from the media (Fig. 6), which demonstrates how a synthetic genetic effect can occur under specific conditions even without drug exposure.

Deletion of *CIT2* induces severe growth defects in *mdj1Δ*. In respiratory strains, *CIT2* deletion does not have the striking effect that *IDH1* or *IDH2* deletion has; however, it does induce greater growth defects under lysine deprivation conditions. TORC1 negatively regulates the RTG pathway through Mks1 [30, 31], which is also involved in the lysine biosynthetic pathway [32]. Constitutive activation of the RTG pathway (via *mks1Δ*) leads to an increase in the expression of lysine biosynthesis enzymes [33]. Thus, the decrease in RTG-target genes under lysine deprivation may coincide with a decrease in lysine biosynthesis enzymes, possibly through a TOR-regulated pathway.

Finally, we see that in WT and all tested HSP mutants, canavanine reliably decreases RTG-target gene expression (Fig. 2B). Even WT shows growth defects in only 2 µg/ml canavanine (Fig. 4).

The combination of 2 µg/ml canavanine with heat shock under arginine deprivation is synthetically lethal for both *cpr7Δ* and *hsp78Δ*, and also inhibits growth in WT and *sse2Δ*. Although expression data shows canavanine strongly reduces RTG-target gene expression in *sse2Δ*, this particular mutant has respiration capacity, no known mitochondrial defects, and shows a robust RTG response under arginine deprivation; thus, it can possibly maintain sufficient expression of RTG-target genes even under canavanine-induced inhibition of the RTG pathway.

Thialysine has a differential effect in HSP mutant backgrounds - some genes are strongly upregulated, some are considerably downregulated, and some RTG-target genes show little response at all in specific backgrounds. Of note, *mdj1Δ* shows a mild change in RTG-target expression in response to thialysine; it’s unclear if this can explain its tolerance to thialysine, is a case of unresponsiveness, or both. *hsp78Δ* shows a robust upregulation of RTG-target genes under thialysine exposure, but is sensitive to the combination of 3 µg/ml thialysine and heat shock, whereas *sse2Δ*, which shows a variable and muted RTG-target gene response to thialysine, is not.

When we consider all of the data, *IDH1* and *IDH2* have a recurring role under various stress conditions in both WT and protein misfolding mutants - canavanine and thialysine exposure in WT as well as arginine deprivation in HSP mutants induce *IDH1/IDH2* mRNA expression; *hsp78Δidh2Δ*, *sse2Δidh1Δ,* and *sse2Δidh2Δ* cannot survive the combined stress of heat shock and canavanine exposure; and the pronounced effect lysine deprivation has on growth of liquid cultures of respiratory HSP/IDH2 double mutants. Our results show that even in WT without additional stressors, *IDH1* and *IDH2* play a major role in overcoming both arginine and lysine deprivation, which could mean that mitochondrial isocitrate dehydrogenase plays a role in coping with high levels of proteotoxic stress. Current research shows that arginine deprivation combined with canavanine is a promising therapy for solid tumors. Our results suggest that combining certain therapies with lysine deprivation may be more suitable in treating aqueous cancer cells.

Taken together, our findings are supported by recent work showing that the RTG pathway is required to both communicate as well as overcome ER protein misfolding stress induced by tunicamycin [15]. More experiments involving a greater number of HSP mutants, combined with other RTG-target gene mutants (like *PDH1*), as well as TCA-cycle mutants not under RTG control, are needed to fully understand the connection between proteotoxic stress, specific amino acid deficiencies, and TCA-cycle metabolite perturbations.

